# Vitamin C – protective role in oxidative stress conditions induced in human normal colon cells by label free Raman spectroscopy and imaging

**DOI:** 10.1101/2021.04.12.439406

**Authors:** K. Beton, B. Brozek-Pluska

## Abstract

Colorectal cancer is the second most frequently diagnosed cancer worldwide. Conventional diagnostics methods of colorectal cancer, can detect it in advanced stage.

Spectroscopic methods, including Raman spectroscopy and imaging, are becoming more and more popular in medical applications, and allow fast, precise and unambiguous differentiation of healthy and cancerous samples. the most important advantage of Raman spectroscopy is ability to identify biomarkers that help in differentiation of healthy and cancerous cells based on biochemistry of sample and spectra typical for: lipids, proteins, DNA.

The aim of the study was to evaluate the biochemical and structural features of human colon cell lines based on Raman spectroscopy and imaging: normal cells CCD-18 Co, normal cells CCD-18 Co under oxidative stress conditions, normal cells CCD-18 Co at first treated by using tert-Butyl hydroperoxide and then supplemented by vitamin C in high concentration to show the protective role of vitamin C in micromolar concentrations against ROS by spectroscopic methods. Raman data obtained for normal cells injured by ROS were compared with spectra typical for cancerous cells.

Statistically assisted analysis has shown that normal, ROS injured and cancerous human colon cells can be distinguished based on their unique vibrational properties.

The research carried out proves that label-free Raman spectroscopy may play an important role in clinical diagnostics differentiation of normal and cancerous colon cells and may be a source of intraoperative information supporting histopathological analysis.

## Introduction

Colorectal cancer (CRC), the third most common cancer in men and the second in woman worldwide, is the third leading cause of death in the World. The mortality rate from this type of cancer is approximately 60% in the United States and Europe [1]. CRC is characterized with high metastasis and poor prognosis [2–4]. In general, the risk factors for colorectal cancer can be divided into three main groups:

- environmental (e.g. high-fat diet, high-calorie diet, diet low in silage, vegetables and fruit),
- internal (e.g. adenomas, ulcers, Crohn’s syndrome),
- genetic (e.g. familial adenomatous polyposis) [5].

75–95% of CRC cases occur in people without any genetic load [6,7]. Those with a family load for two or more first-degree relatives (such as a parent or sibling) have a two to three fold greater risk to suffer CRC, this group accounts for about 10% of all cases [8,9].The additional risk factors influencing the development of CRC are largely related to aging, male sex [7] and lifestyle including high intake of fat, sugar, alcohol [10], red meat, processed meats, obesity, smoking, and a lack of or insufficient physical activity [6,11,12].

Basically, cancer development is a complex multi-stage process that begins anywhere in the body as cells transform from normal to pathological. These changes may occur on their own or they may be induced by the presence or coexistence of factors that we divide into 3 main groups:

- physical, such as ultraviolet and ionizing radiation;
- chemicals, such as asbestos, tobacco smoke, aflatoxins and arsenic;
- biological, e.g. infections due to viruses, bacteria or parasites.

In the first stage of CRC development healthy cells in the lining of the colon or rectum change and grow, and divide uncontrollably to form a mass called a tumor. Both genetic and environmental factors can change the dynamic of this process. CRC most often begins with a polyp, a non-cancerous growth that can develop on the inner wall of the colon or rectum with increasing age and then can transform into cancer or metastatic cancer [13]. Polyps are protrusions of the mucosa lining the walls of the large intestine that extend beyond its surface, thereby reducing the lumen of the colon.

Each day the content of the human colon can be described as a diverse mix of bile, mucus, gut microflora, fermentation products, unabsorbed food, and products of metabolism, including toxins, mutagens, and dissolved gases. In such an environment, the mucosa of the colon is constantly exposed to dietary oxidants and a variety of bacteria. Permanent exposure of the mucosa and the organism itself to unfavourable conditions may lead to uncontrolled oxidative stress and DNA damage, which may lead to the development of cancer disease. Almost 95% of CRC are glandular carcinomas, while the remaining 5% are squamous, mixed or undifferentiated cell types [1].

The protocols to prevent the occurrence and development of CRC include tests ranging from fecal occult blood tests (iFOBT, gFOBT), blood test, colonoscopy examination of high-risk individuals to detect and remove precancerous lesions using biopsy, molecular research, to drug chemotherapy [9,14]. One of the most popular diagnostic imaging methods for CRC is colonoscopy, which allows to monitor the interior of the colon, detect polypic and cancerous changes in it, and take a biopsy sample. The supporting techniques are colonography (virtual colonoscopy) or X-ray examination. Unfortunately, the colonoscopy does not provide answer about origin of tissue abnormality. The suspicious lesion must be removed and further analyzed by trained pathologist by using the gold standard staining protocols that are particularly time consuming, subjective, and high skills personnel demanding. Moreover, colonic perforation occurring in approximately 1 in 1000 cases of colonoscopy, hemorrhagic complications may also occur during or after examination. When combined with anesthesia, other difficulties may include cardiovascular complications such as a temporary drop in blood pressure, the risk of blood clots, pulmonary embolism or deep vein thrombosis. That’s why the development of new diagnostics methods including those based on Raman spectroscopy and imaging stimulate so many research group worldwide.

The first in-vivo Raman measurements of human gastrointestinal tissue were published in 2000 by Shim et al. [15]. This study has shown that fiber-optic-coupled Raman spectroscopy can be successfully used for disease classification during in-vivo measurements. FT-Raman studies of colorectal cancers have been published also by Andrande et al. [16]. Authors described a diagnostic algorithm useful to establish the spectral differences of the complex colon tissues to find a characteristic Raman features. The first Raman and CARS data of colon tissue were published by Krafft et al. [17]. Authors showed CARS images, whose were recorded from thin colon tissue sections at 2850, 1660, 1450 and 1000 cm^-1^ and compared with Raman images obtained using classical spontaneous scattering Raman effect. Comparison between CARS and spontaneous Raman images confirmed that results obtained using both methods are comparable, but a time needed for CARS maps acquisition is three orders of magnitude shorter. In 2013 Gerwert et al [18] have shown that the auto-fluorescence of colon tissue overlaps spatially with the fluorescence of antibodies against p53, which are of interest in routine immunohistochemistry in pathology analysis and indicates nuclei with mutated p53 of cancer cells. They have also shown many advantages of VIS region (532 nm) excitation compared to excitations wavelengths from IR region (785 or 830 nm) most often used in spectroscopic experiments [18].

Despite the continuous development of synthetic anticancer drugs, there is still a need to discover and research natural compounds with a therapeutic effect on the incidence and progression of cancer [19]. Plants and microorganisms are a rich source of molecules that can help fight cancer and are the perfect combination for supporting the natural immune mechanism. Natural anti-cancer drugs do not wreak havoc on the human body and side effects and, when given in the right form, are much better absorbed. In addition, natural ingredients are the substrates of many physiological reactions, and their presence affects the quality and dynamic of processes taking place in the body. An example of such physiologically important compounds are vitamins. In generally, vitamins are organic molecules that are essential for a wide variety of cellular functions as coenzymes, antioxidants and regulators of gene expression. Vitamins available in nature can be divided into fat-soluble vitamins (A, D, E, K) and water-soluble vitamins (B1, B2, B3, B5, B6, B7, B9, B12, C).

Vitamin C (ascorbic acid) is an essential micronutrient that must be provided in the diet or as a supplement. People have lost the ability to synthesize it naturally due to a mutation in the gene encoding the final enzyme in its biosynthetic pathway [20]. Vitamin C plays a significant role in many processes as a cofactor of enzymes involved in processes important for cancer development: antioxidant defence, transcription and epigenetic regulation of gene expression. Vitamin C has also been shown to have a beneficial effect on the immune system by mobilizing it to fight infections, stimulating, for example, the production of white blood cells, and also prevents inflammation, which is key in the body’s fight against cancer cells [21]. The anti-cancer potential of vitamin C is suggested by the results of many laboratory studies in animals and cell cultures [22–30]. In anti-cancer research, not only vitamin C is used, but also its derivatives, including compounds with increased lipophilicity and resistance to oxidation [31].

Vitamins are also the class of molecules that exert an antioxidant effect either directly through an internal free radical scavenging mechanism or indirectly by participating in the regulation and expression of enzymes (e.g. inducible nitric oxide synthesis (iNOS)).

Vitamin C is the most important antioxidant representing the group of water-soluble antioxidants. In principle, the action of vitamin C takes place by direct scavenging of ROS, especially peroxides and peroxynitrite [32]. Moreover, in the work of Diaz et al. [33] has been shown that ascorbate can revitalize alpha-tocopherol, which in turn helps prevent lipid oxidation in the body. On the other hand, without ascorbate, the alpha-tocopheroxyl radical may play a pro-oxidative role and continue or even enhance the lipid peroxidation process.

By acting as an electron donor, vitamin C can also reduce superoxide anions, hydroxyl radicals, singlet oxygen, and hypochlorous acid generated during metabolic respiration where ATP is produced. Vitamin C can therefore protect against mutations caused by oxidative DNA damage, lipid peroxidation and oxidation of amino acid residues in order to maintain protein integrity by inhibiting the emission of free radicals [34].

The level of vitamin C in human plasma is tightly regulated to maintain a physiological concentration of about 50-70 µM, which limits its ability to be highly concentrated in cells just by oral ingestion. As it is well known, oral consumption of an agent with a given concentration does not make its concentration in a given organ inside the body identical. The oral route of ingestion of e.g. vitamin C is the longest possible route. When vitamin C is ingested, it is absorbed from the colon lumen and released into the bloodstream. In the digestive tract, the ionized form of vitamin C ascorbate (ASC) and its oxidized counterpart, dehydroascorbic acid (DHA), are absorbed by various transport systems. Distribution from plasma to tissue is regulated differently, and organs and tissues differ significantly in their vitamin C content. The tight regulation of vitamin C homeostasis is primarily controlled by four regulatory systems [35]:

- intestinal absorption,
- accumulation and distribution in tissues,
- degree of use and recyclability for reuse,
- renal excretion and reabsorption.

This can be accomplished through a variety of mechanisms, including passive diffusion, facilitated diffusion, active transport, and re-use through a reabsorption system [36]. For the above reasons, scientific research on antioxidant delivery conducted using the intravenous method, omitting the so-called first-pass effect associated with oral supplementation, has gained popularity.

The first documented studies of high-dose intravenous vitamin C in cancer therapy were published in the 1970s [22]. A few years later, clinical data showed that when ascorbate is administered orally, its concentration in plasma is tightly controlled by other organs and enzyme systems ensuring active absorption [37]. After oral administration of vitamin C, its steady state plasma concentration is approximately 2,500 times lower. When doses exceed 200 mg, relative absorption decreases, urinary excretion increases, and the proportion of bioavailable ascorbate decreases [38]. Maximum values in plasma do not exceed ≈ 220 µM even after a maximum oral dose of 3g 6 times a day [39]. In contrast, when ascorbate is administered intravenously, millimolar concentrations can be achieved. Thus, many independent scientific reports confirm that only intravenous administration of ascorbate can cause pharmacological, therapeutic levels of the antioxidant [27,39,40].

Vitamin C at normal physiological concentrations acts as a water-soluble antioxidant. However, at high concentrations (350-450 mg/dl), vitamin C dissociates in the extracellular fluid and transforms into an ascorbate radical (AscH-), reducing the iron oxidation state from + III to + II [27]. The reduced iron then reacts with the oxygen to form the superoxide anion (O_2_^-^), which reacts with the hydrogen to form H_2_O_2_. The increased concentration of H_2_O_2_ in cancerous cells makes the cells vulnerable to the cytotoxic effect of H_2_O_2_ [27,28,41]. The consequence of this is the apoptosis of cells, which leads to their death. At the same time, the high concentration of vitamin C in the serum inhibits the reduction of glutathione, causing its presence in the oxidized form. As a result of this reaction, H_2_O_2_ accumulates, which in normal, healthy cells is easily metabolized to oxygen and water by the enzyme catalase. On the other hand, cancer cells lack the catalase enzyme or its action is defective. This process of selective cytolytic activity makes it possible to effectively target cancer cells without affecting the body’s immune cells [27]. In addition to the absence of catalase, cancer cells selectively take up more vitamin C compared to normal cells due to the increased regulation of glucose transporters to meet their metabolic needs.

In scientific literature [28,29] it was shown that some cancerous cells, compared to normal cells, have an increased sensitivity to ascorbate-induced cytotoxicity. The difference in the sensitivity of normal and cancerous cells to ascorbate may be due to the low level of antioxidant enzymes and the high level of endogenous ROS in cancerous cells [42,43]. The relatively lower activity of catalase, glutathione peroxidase, and peroxyredoxins in cancer cells could potentially contribute to less effective H_2_O_2_ removal and increased sensitivity to ascorbate-induced cytotoxicity.

Free radicals, called reactive oxygen species (ROS), are constantly produced in animal and human cells. Excess ROS can cause oxidative damage to cells and promote many degenerative diseases as well as accelerate the aging process due to the formation of oxidative stress in the body.

Under normal, balanced conditions, the body itself copes with the elimination of reactive oxygen species through antioxidants supplied to it with food. However, when the body is in a stressful, longer-lasting situation or when we lead an unhealthy lifestyle, the concentration of free radicals can increase significantly, and the body will find it difficult to neutralize them. Chronic oxidative stress is the initiating stage of many health problems, as the excess of reactive oxygen species contributes to the intensification of destructive processes. One of them is the oxidation of structural proteins, lipids and cell membranes. Permanent, high concentration of reactive oxygen species also leads to DNA damage, which may cause mutations in the body, gene expression blockages, impaired immune reactions, and eventually cancer of virtually any organ can develop. Simultaneously ROS are essential for the proper conduct of a number of protective reactions. They are the mediators of antimicrobial phagocytosis, detoxification reactions led by the cytochrome P-450 complex and apoptosis.

Providing the body with exogenous antioxidants such as vitamin C, vitamin E and beta-carotene may protect against cancer and other diseases in people with congenital or acquired high levels of ROS [44].

The aim of the study was to evaluate the biochemical features of human colon cell lines based on label-free Raman spectroscopy and imaging: normal CCD-18 Co, normal CCD-18 Co under oxidative stress conditions, normal CCD-18 Co at first treated by using tert-Butyl hydroperoxide and then supplemented by vitamin C in micromolar concentration to show the protective role of vitamin C against Reactive Oxygen Species. Data attained for normal cells injured by ROS were compared and contrasted with data typical for cancerous cells CaCo-2.

Statistically assisted analysis of Raman data shows that normal, ROS injured and cancerous human cells of human colon can be distinguished based on their unique vibrational properties typical for proteins, lipids and DNA.

The research carried out proves that label-free Raman spectroscopy may play an important role in clinical diagnostics and in the future may be a source of intraoperative information supporting histopathological analysis in differentiation of healthy and cancer cells.

## Materials and methods

### Cell lines and cell culture

CCD-18Co cell line (ATCC^®^ CRL-1459™) was purchased from ATCC: The Global Bioresourece Center. CCD-18Co cell line was cultured using ATCC-formulated Eagle’s Minimum Essential Medium with L-glutamine (catalog No. 30-2003). To make the complete growth medium, fetal bovine serum was added to a final concentration of 10%. Every 2–3 days, a new medium was used. The cells obtained from the patient are normal myofibroblasts in the colon. The biological safety of the CCD-18Co cell line has been classified by the American Biosafety Association (ABSA) as level 1 (BSL-1). The CaCo-2 cell line was also purchased from ATCC and cultured according to the ATCC protocols. The CaCo-2 cell line was obtained from a patient - a 72-year-old Caucasian male diagnosed with colon adenocarcinoma. The biological safety of the obtained material is classified as level 1 (BSL - 1). To make the medium complete we based on Eagle’s Minimum Essential Medium with L-glutamine, with addition of a fetal bovine serum to a final concentration of 20%. The medium was renewed once or twice a week.

### Cultivation conditions

Cell lines (CCD-18Co, Caco-2) used in the experiments in this study were grown in flat-bottom culture flasks made of polystyrene with a cell growth surface of 75 cm^2^. Flasks containing cells were stored in an incubator providing environmental conditions at 37 °C, 5% CO_2_, 95% air.

### Cell treatment with vitamin C and/or tBuOOH

Cells used for research were seeded onto CaF_2_ windows (25 × 1 mm) at a low density of 10^4^ cells/cm^3^. After 24 h incubation on the CaF_2_, standard growth medium was removed and vitamin C solution was added for 24 or 48 hours. For the stress agent variant, tBuOOH at a concentration of 50 µM with or without 50 µM of vitamin C was added. After 24 or 48 hours, the cells were rinsed with phosphate-buffered saline (PBS, Gibco, 10010023, pH 7.4 at 25°C, 0.01 M) to remove any residual medium and an excess vitamin C that did not penetrate inside the cells. Furthermore, PBS was removed and cells were fixed in paraformaldehyde (4% buffered formaldehyde) for 10 min, and washed once more with PBS. The Raman confocal measurements were made immediately after the preparation of the samples.

All the stress compound solutions, based on tBuOOH, were prepared by pre-diluting in the PBS solvent. After obtaining the initial concentration of 20,000 μM, it was diluted in a medium intended for a given type of cells, so as to obtain the concentration of 200 μM and finally 50 μM.

### Raman Spectroscopy and Imaging

All maps and Raman spectra presented and discussed in this paper were recorded using the confocal microscope Alpha 300 RSA+ (WITec, Ulm, Germany) equipped with an Olympus microscope integrated with a fiber with 50 µm core diameter with a UHTS spectrometer (Ultra High Through Spectrometer) and a CCD Andor Newton DU970NUVB-353 camera operating in default mode at -60 °C in full vertical binning mode. 532 nm excitation laser line, which is the second harmonic of the Nd: YAG laser, was focused on the sample through a Nikon objective lens with magnification of 40x and a numerical aperture (NA = 1.0) intended for cell measurements performed by immersion in PBS. The average excitation power of the laser during the experiments was 10 mW, with an integration time of 0.5 s for Raman measurements for the high frequency region and 1.0 s for the low frequency region. An edge filter was used to filter out the Rayleigh scattered light. A piezoelectric table was applied to set the test sample in the right place by manipulating the XYZ positions and consequently record Raman images. Spectra were acquired with one acquisition per pixel and a diffraction grating of 1200 lines/mm. Cosmic rays were removed from each Raman spectrum (model: filter size: 2, dynamic factor: 10) and the Savitzky-Golay method was implemented for the smoothing procedure (order: 4, derivative: 0). All data was collected and processed using a special original software WITec Project Plus. All imaging data were analyzed by Cluster Analysis (CA), which allows for grouping of a set of vibrational spectra that bear resemblance to each other. CA was executed using WITec Project Plus software with Centroid model and k-means algorithm, in which each cluster is represented by one vector of the mean. Data normalization was performed using the model: divided by norm, which was wrought with the Origin software serving as mathematical and statistical analysis tool.

### Chemical compounds

L-Ascorbic acid, reagent grade, crystalline catalogue Number A7506-25G, Luperox^®^ TBH70X, tert-Butyl hydroperoxide solution catalogue Number 458139, bisBenzimide H 33342 trihydrochloride catalogue Number B2261, Red Oil-O catalogue Number O0625 were purchased from Sigma-Aldrich, and used without additional purification. XTT proliferation Kit with catalogue Number 20-300-1000 was purchased from Biological Industries.

### XTT cells viability tests

The XTT test is grounded on the cleaving of the yellow tetrazolium salt XTT to form an orange water soluble formazan product by activity of dehydrogenase enzyme in the active mitochondria. The quantity of formazan product is directly proportional to the number of living and respiring cells. Diminution in amount of living cells results in a depletion in the overall activity of mitochondrial dehydrogenases in the sample. This decrease directly is in consonance with the amount of the generated orange formazan, monitored by the measurement of absorbance using multi-sensing microplate spectrophotometer. With the XTT test, cell proliferation and viability of the cells after treatment with the test item are determined colorimetrically. The use of the XTT reagent consist in the statistical calculation of the metabolic activity of cells wherewithal a colorimetric techniques in response to changed environmental conditions in which they are located.

The XTT test requires the usage of a 450 nm wavelength, from which the specific signal of the sample is obtained, and 650 nm, which is the referential sample during the test. XTT reagent is used to computing the activity of metabolic processes in living cells. Tetrazolium salts are converted in cells by means of special enzymes to formazan, however, this reaction takes place properly only in cells with undamaged metabolism. XTT is believed to be excluded from entering cells due to a negative net charge [45]. Scientific work suggests that the reduction of the XTT dye transpire at the cell surface, which facilitates the transport of electrons across the cytoplasmic membrane. Mitochondrial oxidoreductases are believed to contribute significantly to the response to the XTT reagent and their reducers are transferred to the plasmalemma. Ultimately, it was found that XTT assays actually measure the redox state of pyridine nucleotide cells [45,46].XTT tests on cell lines were carried out on a multi-detecting BioTek Synergy HT model reader designed for microplate testing. The test protocol was created especially for the presented experiments.

Scheme 1 shows the results of XTT test obtained for CCD-18Co human normal colon cells supplemented with Vitamin C in various concentrations and in various time intervals.

**Scheme 1:**
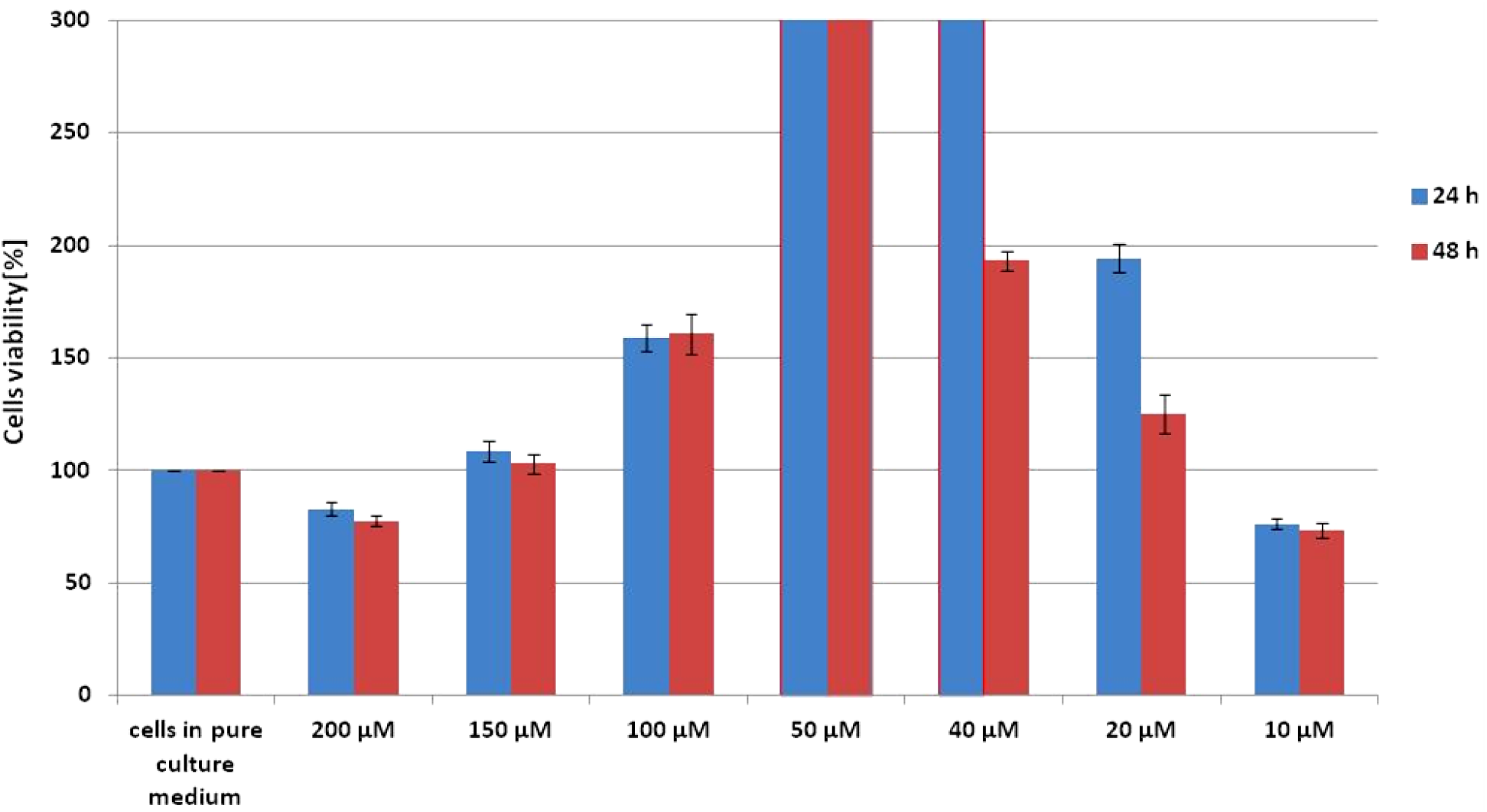
Results of XTT comparison of the percent viability for CCD-18Co human normal colon cells supplemented with different concentrations of Vitamin C in two different time intervals with the standard deviation ±SD.

The addition of vitamin C at different concentrations to cells with a normal structure does not cause damage in the cells, which can be concluded from the fact that their viability rate is higher than for the referential sample. For the highest and the lowest concentration of vitamin C, a slight decrease in survival below 100% can be observed. The attached data show that the most optimal concentration for this cell line, the action of which stimulate the metabolic processes of cells, is 50 μM. The basis for such a conclusion is the percentage of cell viability, which for this concentration is above the measuring range of the device, allows us to conclude that cells in such an environment grow and multiply more often and faster. This result is identical for incubation times of 24 and 48 hours, so it can be suspected that maintaining a constant saturation of colon cells with vitamin C at a concentration of 50 μM supports the proper proliferation and vital functions of cells.

## Results

In this section data obtained by Raman spectroscopy and imaging for human colon cells both before and after the generation of Reactive Oxygen Species (ROS), including supplementation with vitamin C are expounded. We will also present a comparison of spectra for the normal human colon cells (CCD-18 Co), for normal colon cells under oxidative stress conditions and for cancerous human colon cell line (CaCo-2).

Generally, Raman vibrational spectra consists of two interesting regions: the Raman fingerprint region: 500–1800 cm^-1^ and the high frequency region: 2700–3100 cm^-1^ (the region 1800-2700 cm^-1^ is excluded from consideration through the lack of Raman bands).

To properly rise to biochemical changes in both normal and cancerous human colon cell lines by Raman spectroscopy and imaging, we will closely inquire how they Raman based method responds to generated ROS. The experiments will extend our knowledge of the protective effect (function) of antioxidants and veritable influence of the ROS generation on cancer development.

Figure 1 presents the microscopy image, Raman image of human colon normal single cell CCD-18Co constructed based on Cluster Analysis (CA) method, Raman images of all clusters identified by CA assigned to: lipid-rich regions, mitochondria, nucleus, cytoplasm, cell membrane, and cell environment, the average Raman spectra typical for CCD-18Co human normal colon cell for all identified clusters for low frequency and high frequency region, and the average Raman spectrum for human normal colon cell - for cells as a whole, all data for experiments performed without any supplementation, cells measured in PBS, colors of the spectra correspond to the colors of clusters.

**Figure 1.**
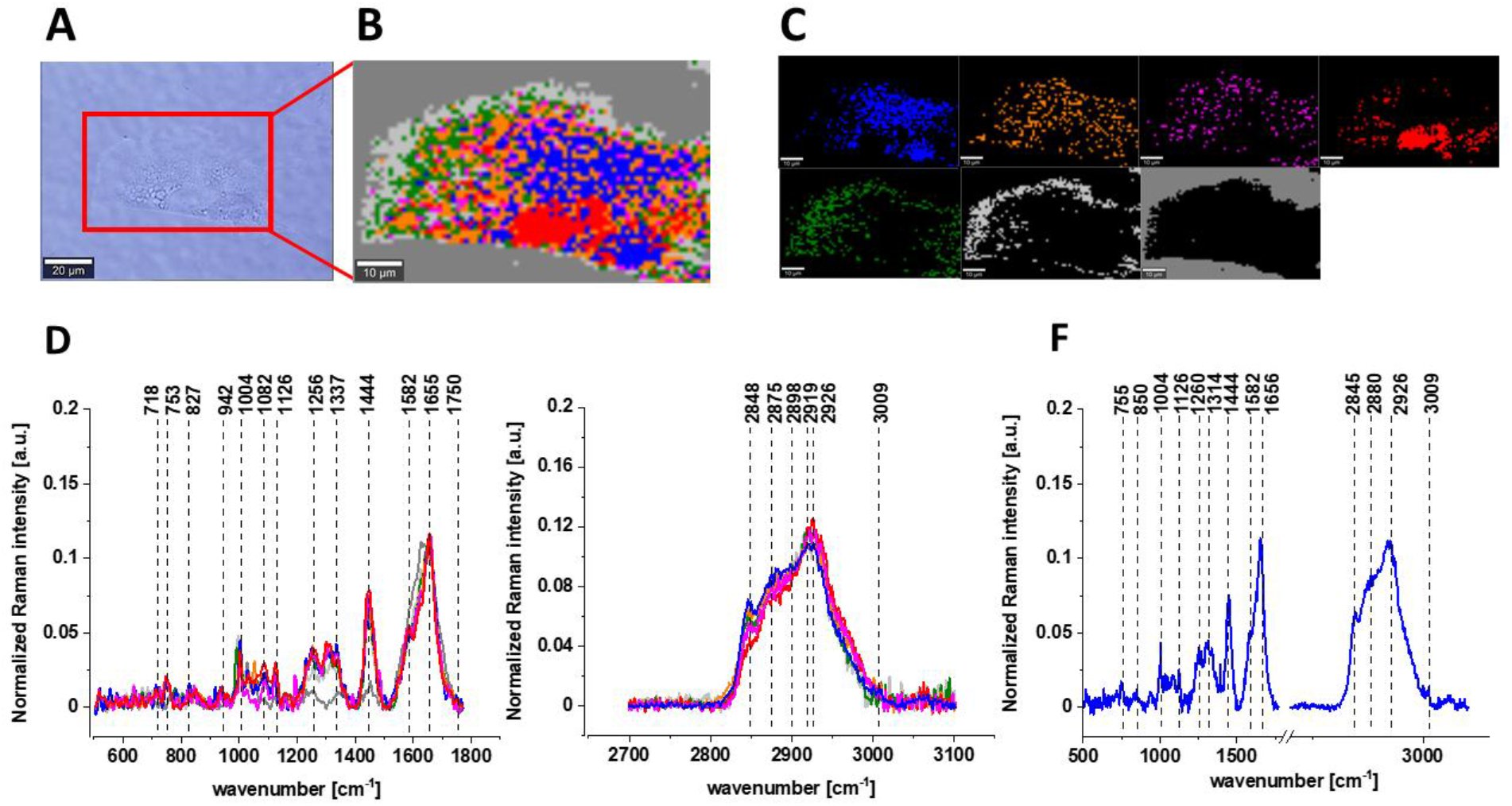
The microscopy image (A), Raman image (B) of human colon normal single cell CCD-18Co constructed based on Cluster Analysis (CA) method, Raman images of all clusters identified by CA assigned to: lipid-rich regions (blue and orange), mitochondria (magenta), nucleus (red), cytoplasm (green), cell membrane (light grey), and cell environment (dark grey) (C), the average Raman spectra typical for CCD-18Co human normal colon cell for all identified clusters for low frequency and high frequency region (D), and the average Raman spectrum for human normal colon cell - for cell as a whole (E), all data for experiments performed without any supplementation, cells measured in PBS, colors of the spectra correspond to the colors of clusters, excitation laser line 532 nm.

As we have mentioned above ROS are produced by living organisms as a result of natural cellular metabolism, but in such a case the concentrations of them is low to moderate functions in physiological cell processes, but at high ROS concentrations adverse modifications to cell components, such as: lipids, proteins, and DNA can be noticed [47]. The shift in balance between oxidant/antioxidant in favour of oxidants seems to be crucial for understanding many dysfunctions of human cells including cancer transformation. But before we start the analysis of ROS treated human normal colon cells let’s focus on CCD-18Co cells supplemented with vitamin C in micromolar concentration to check how vitamin influences the cells vibrational features.

Figure 2 presents the microscopy image, Raman image of human colon normal single cell CCD-18Co after 24h of vitamin C supplementation constructed based on Cluster Analysis (CA) method, Raman images of all clusters identified by CA assigned to: lipid-rich regions, mitochondria, nucleus, cytoplasm, cell membrane, and cell environment, the average Raman spectra typical for CCD-18Co human normal colon cell for all identified clusters for low frequency and high frequency region, and the average Raman spectrum for human normal colon cell - for cells as a whole, all data for experiments performed with supplementation with vitamin C of 50 μM concentration in medium, cells measured in PBS, colors of the spectra correspond to the colors of clusters.

**Figure 2.**
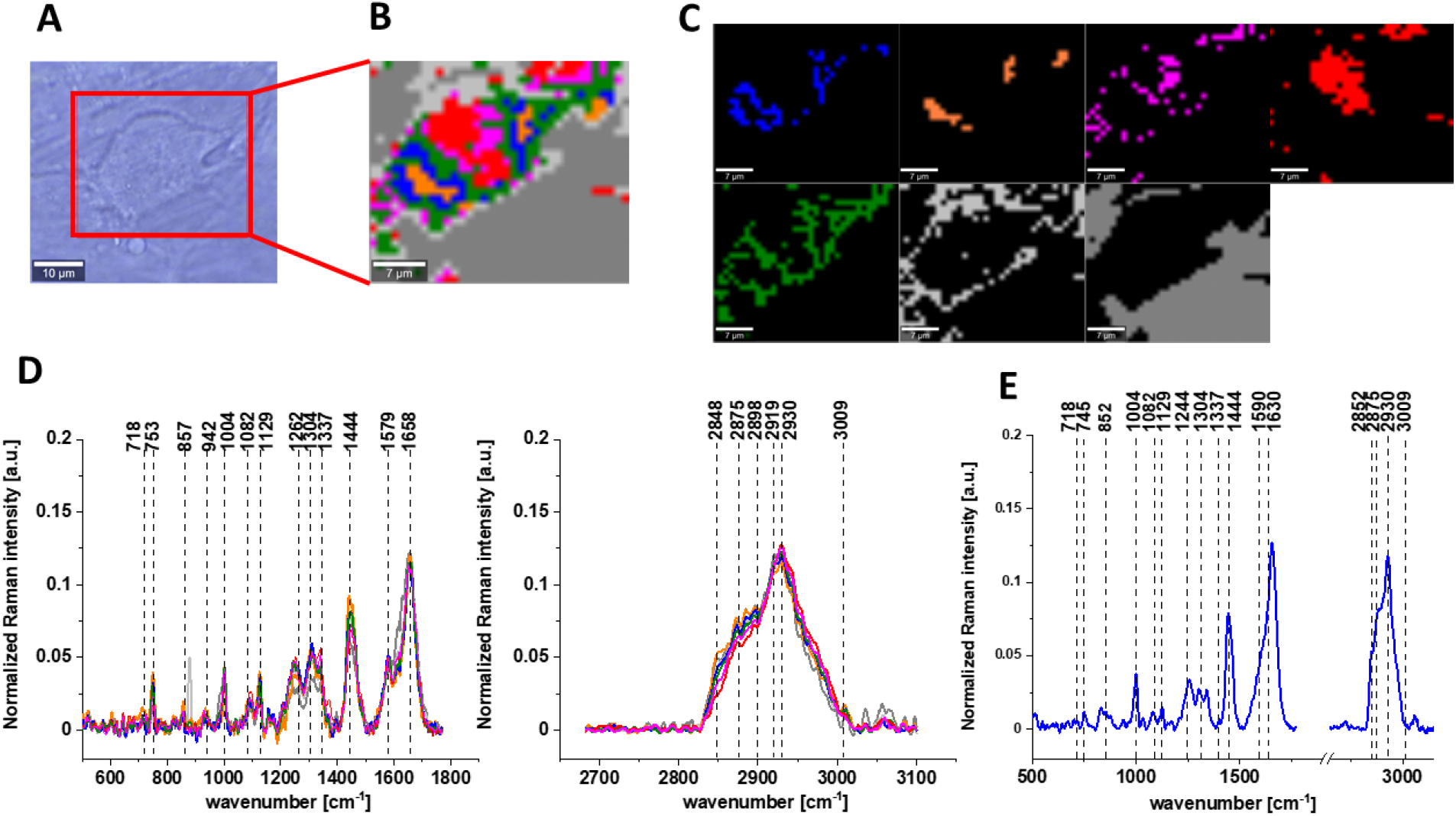
The microscopy image (A), Raman image (B) of human colon normal single cell CCD-18Co after 24h of vitamin C supplementation constructed based on Cluster Analysis (CA) method, Raman images of all clusters identified by CA assigned to: lipid-rich regions (blue and orange), mitochondria (magenta), nucleus (red), cytoplasm (green), cell membrane (light grey), and cell environment (dark grey) (C), the average Raman spectra typical for CCD-18Co human normal colon cell for all identified clusters for low frequency and high frequency region (D), and the average Raman spectrum for human normal colon cell - for cell as a whole (E), cells measured in PBS, colors of the spectra correspond to the colors of clusters, excitation laser line 532 nm.

The same Raman analysis was performed also for human normal colon cells CCD-18Co with supplementation by vitamin C of 50 μM concentration in medium for incubation time 48h. Figure 3 shows the microscopy image, Raman image of human colon normal single cell CCD-18Co after 48h of vitamin C supplementation constructed based on Cluster Analysis (CA) method, Raman images of all clusters identified by CA assigned to: lipid-rich regions, mitochondria, nucleus, cytoplasm, cell membrane, and cell environment, the average Raman spectra typical for CCD-18Co human normal colon cell for all identified clusters for low frequency and high frequency region, and the average Raman spectrum for human normal colon cell - for cell as a whole, all data for experiments performed with supplementation with vitamin C of 50 μM concentration in medium, cells measured in PBS, colors of the spectra correspond to the colors of clusters.

**Figure 3.**
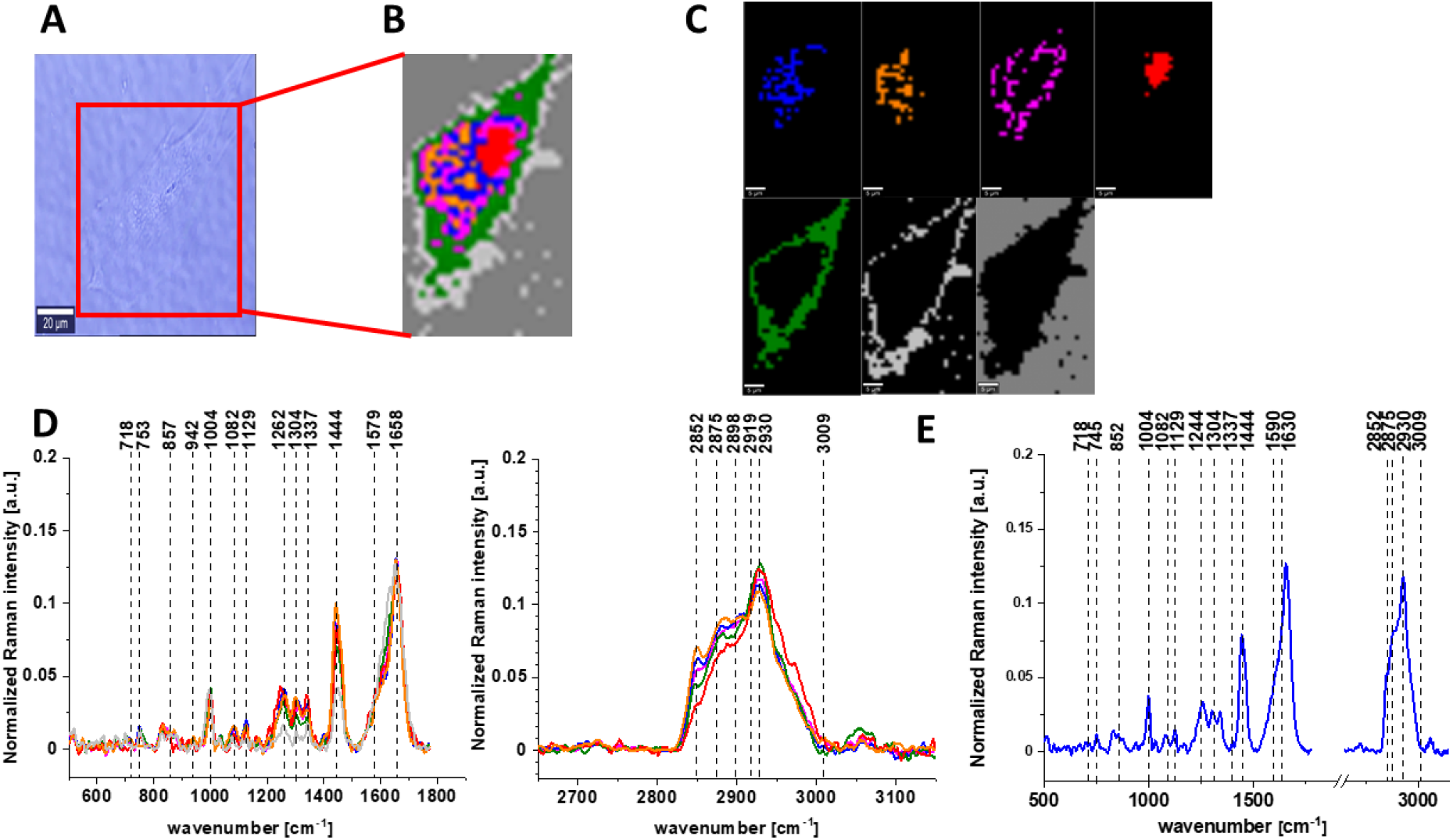
The microscopy image (A), Raman image (B) of human colon normal single cell CCD-18Co after 48h of vitamin C supplementation constructed based on Cluster Analysis (CA) method, Raman images of all clusters identified by CA assigned to: lipid-rich regions (blue and orange), mitochondria (magenta), nucleus (red), cytoplasm (green), cell membrane (light grey), and cell environment (dark grey) (C), the average Raman spectra typical for CCD-18Co human normal colon cell for all identified clusters for low frequency and high frequency region (D), and the average Raman spectrum for human normal colon cell - for cell as a whole (E), cells measured in PBS, colors of the spectra correspond to the colors of clusters, excitation laser line 532 nm.

The next step of our experiments was the analysis of Raman features typical for human colon normal cells CCD-18Co under oxidative stress conditions generated by using tBuOOH on concentration of 50 μM in medium for 24 and 48h.

Figures 4 and 5 show the microscopy images, Raman images of human colon normal single cells CCD-18Co under oxidative stress conditions generated by using tBuOOH on concentration of 50 μM in medium for 24 and 48h respectively.

**Figure 4.**
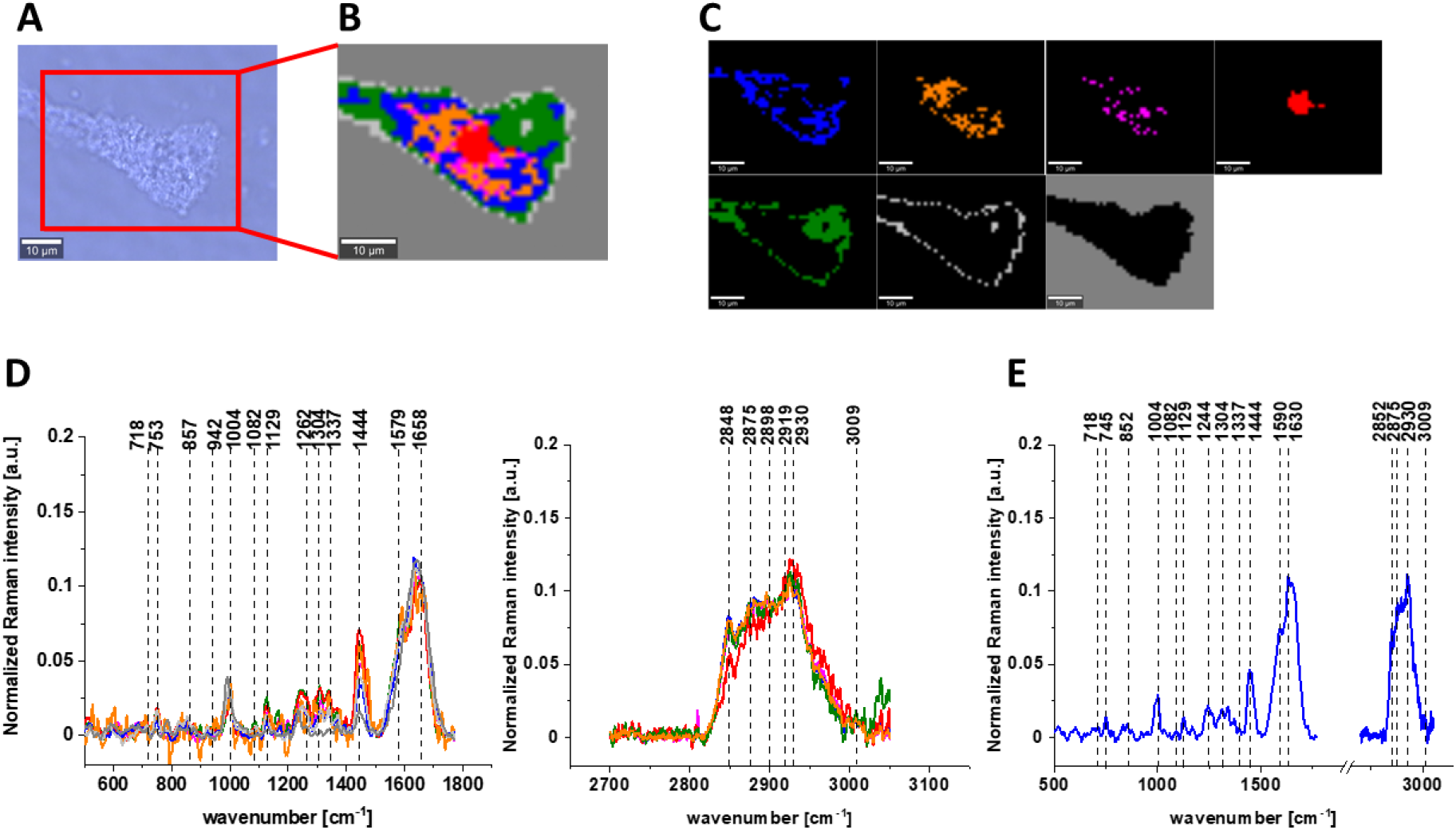
The microscopy image (A), Raman image (B) of human colon normal single cell CCD-18Co under oxidative stress conditions generated by using tBuOOH for incubation time of 24h and concentration 50 μM in medium constructed based on Cluster Analysis (CA) method, Raman images of all clusters identified by CA assigned to: lipid-rich regions (blue and orange), mitochondria (magenta), nucleus (red), cytoplasm (green), cell membrane (light grey), and cell environment (dark grey) (C), the average Raman spectra typical for CCD-18Co human normal colon cell for all identified clusters for low frequency and high frequency region (D), and the average Raman spectrum for human normal colon cell - for cell as a whole (E), cells measured in PBS, colors of the spectra correspond to the colors of clusters, excitation laser line 532 nm.

**Figure 5.**
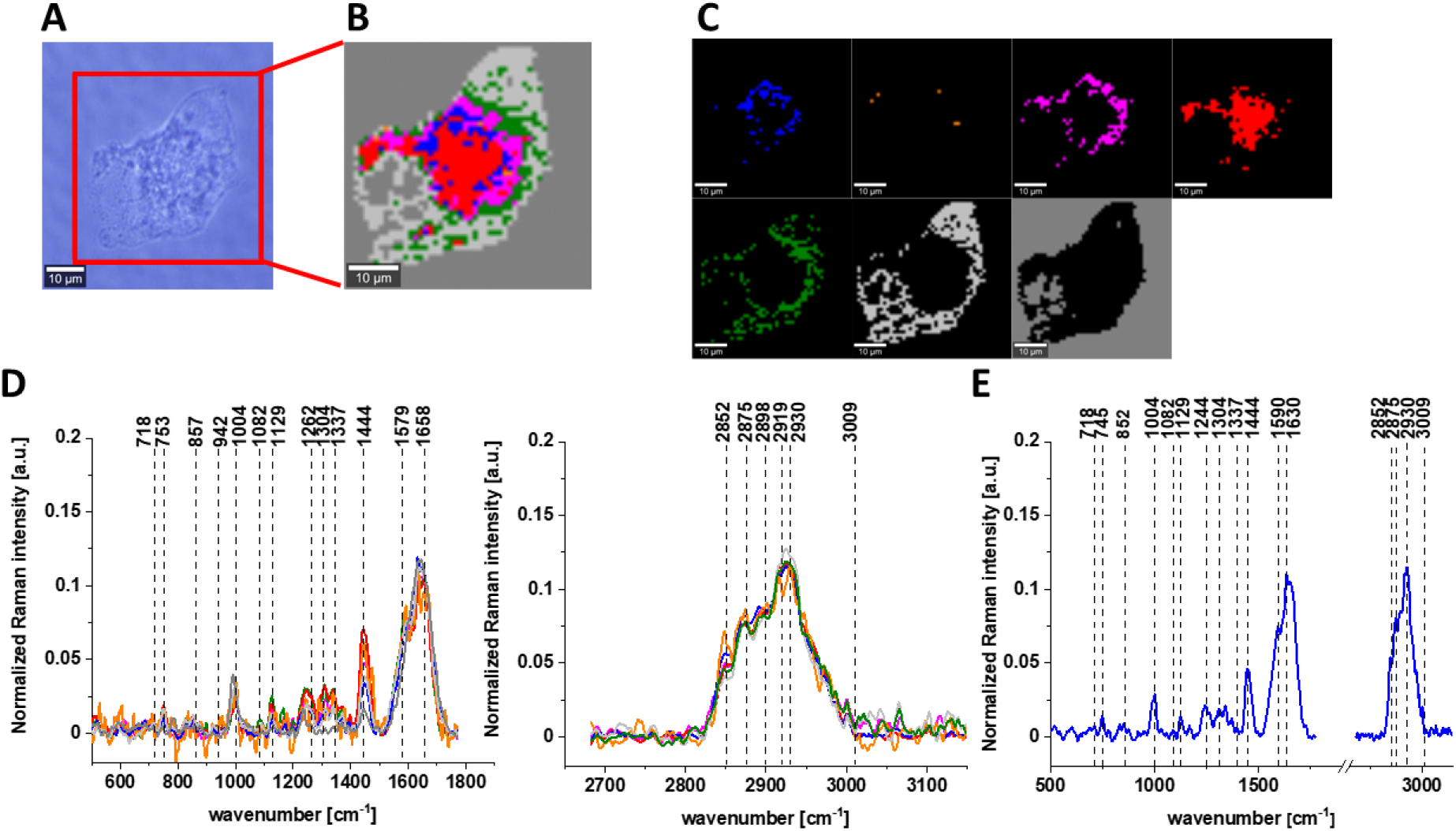
The microscopy image (A), Raman image (B) of human colon normal single cell CCD-18Co under oxidative stress conditions generated by using tBuOOH for incubation time of 48h and concentration 50 μM in medium constructed based on Cluster Analysis (CA) method, Raman images of all clusters identified by CA assigned to: lipid-rich regions (blue and orange), mitochondria (magenta), nucleus (red), cytoplasm (green), cell membrane (light grey), and cell environment (dark grey) (C), the average Raman spectra typical for CCD-18Co human normal colon cell for all identified clusters for low frequency and high frequency region (D), and the average Raman spectrum for human normal colon cell - for cells as a whole (E), cells measured in PBS, colors of the spectra correspond to the colors of clusters, excitation laser line 532 nm.

Next to check if the vitamin C shows the protective properties against ROS we have measured human normal colon cells CCD-18Co supplemented simultaneously by tBuOOH and vitamin C.

Figures 6 and 7 show the vibrational data obtained for human normal cells CCD-18Co for experiments performed with adding of tBuOOH and vitamin C for 24 or 48h by Raman spectroscopy and imaging.

**Figure 6.**
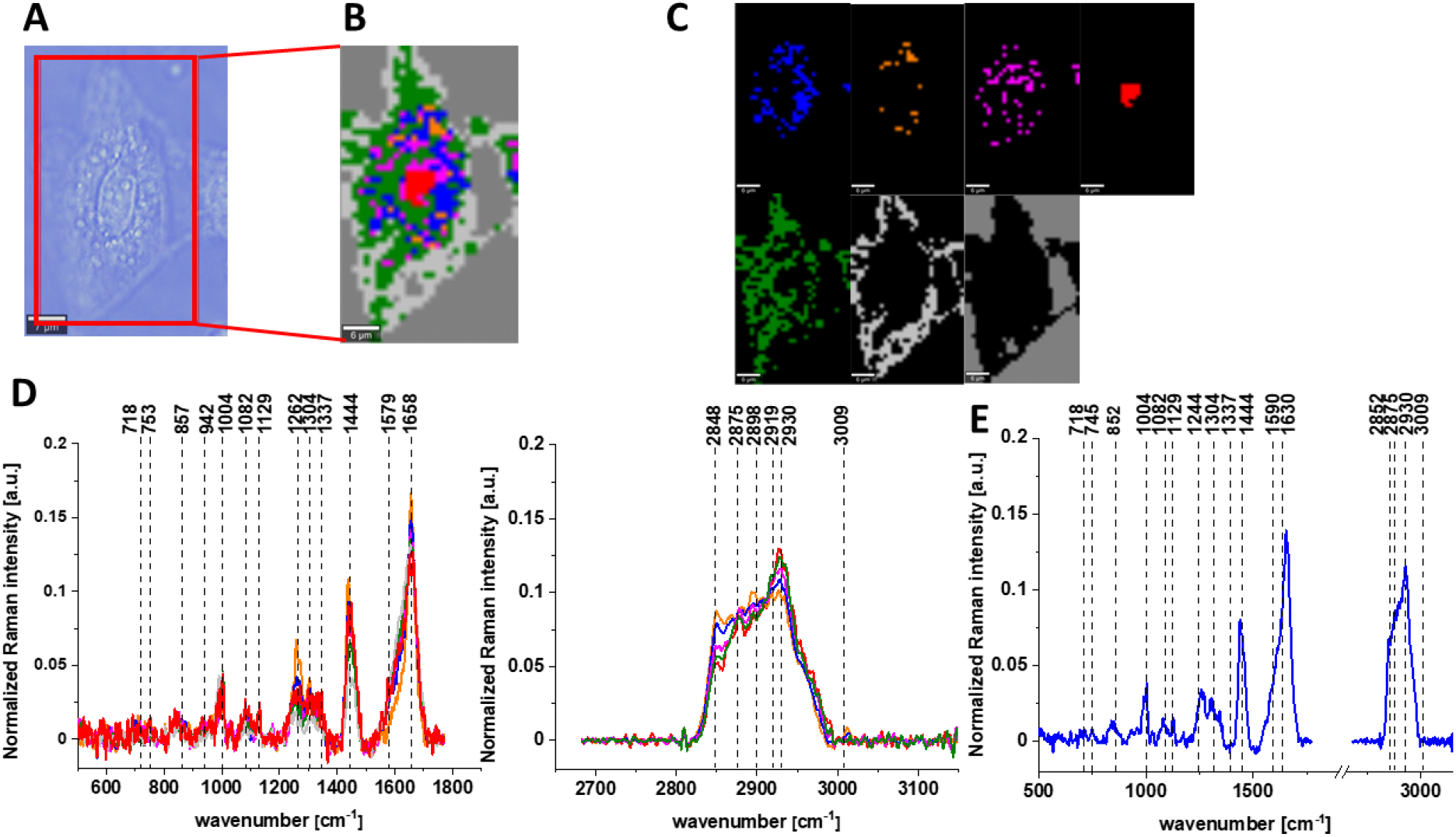
The microscopy image (A), Raman image (B) of human colon normal single cell CCD-18Co for supplementation by tBuOOH (50 μM in medium) and vitamin C (50 μM in medium) for incubation time of 24h constructed based on Cluster Analysis (CA) method, Raman images of all clusters identified by CA assigned to: lipid-rich regions (blue and orange), mitochondria (magenta), nucleus (red), cytoplasm (green), cell membrane (light grey), and cell environment (dark grey) (C), the average Raman spectra typical for CCD-18Co human normal colon cell for all identified clusters for low frequency and high frequency region (D), and the average Raman spectrum for human normal colon cell - for cell as a whole (E), cells measured in PBS, colors of the spectra correspond to the colors of clusters, excitation laser line 532 nm.

**Figure 7.**
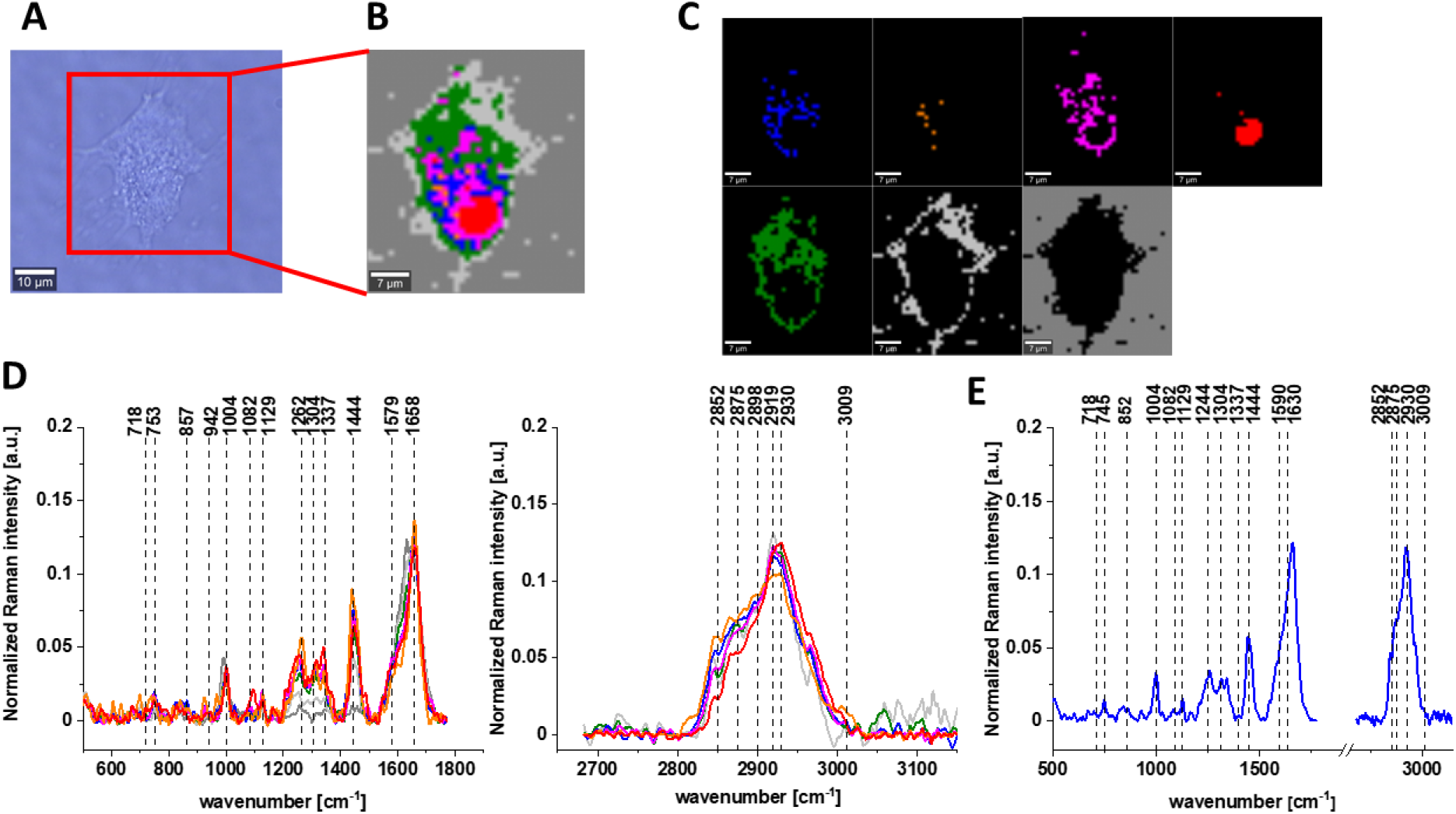
The microscopy image (A), Raman image (B) of human colon normal single cell CCD-18Co for supplementation by tBuOOH (50 μM in medium) and vitamin C (50 μM in medium) for incubation time of 48h constructed based on Cluster Analysis (CA) method, Raman images of all clusters identified by CA assigned to: lipid-rich regions (blue and orange), mitochondria (magenta), nucleus (red), cytoplasm (green), cell membrane (light grey), and cell environment (dark grey) (C), the average Raman spectra typical for CCD-18Co human normal colon cell for all identified clusters for low frequency and high frequency region (D), and the average Raman spectrum for human normal colon cell - for cell as a whole (E), cells measured in PBS, colors of the spectra correspond to the colors of clusters, excitation laser line 532 nm.

Table 1 featured the main chemical constituents which can be identified based on their vibrational properties in analyzed CCD-18Co human, normal colon cells in normal conditions, in oxidative stress conditions generated by tBuOOH adding (discussed later in the manuscript) and for the experiments for vitamin C supplementation.

**Table 1.** Band positions and tentative assignments for human normal colon cells from control sample, in oxidative stress conditions generated by tBuOOH adding (discussed later in the manuscript) and for the experiments for vitamin C supplementation. Data based on the average Raman spectra for cells as a whole measured in PBS [48].

| Wavelength [$\text{cm}^{-1}$] | Tentative assignments |
| --- | --- |
| 755 | Nucleic acids, DNA, Tryptophan, Nucleoproteins |
| 850 | Tyrosine |
| 1004 | Phenylalanine |
| 1126 | Saturated fatty acids, cytochrome C |
| 1260 | Amide III (C–N stretching + N–H bending) |
| 1314 | $\text{CH}_3\text{CH}_2$ twisting mode of collagen |
| 1444 | Lipids and Proteins |
| 1582 | $\text{CN}_2$ scissoring and $\text{NH}_2$ rocking of mitochondria and |
|  | phosphorylated proteins |
| 1656 | Amide I (C=O stretch) |
| 2848 | Lipids, fatty acids |
| 2880 | Lipids and proteins |
| 2931 | Due primarily to protein |
| 3009 | (=C-H), lipids, fatty acids |

The main goal of the research undertaken in this study was the biochemical analysis of human normal colon cell line in normal or ROS conditions, demonstrating the antioxidant properties of vitamin C and comparative analysis of normal human colon cells in different conditions with human cancerous colon cells CaCo-2 based on the vibrational spectra of colon cells using label-free Raman spectroscopy and imaging. Figure 8 shows Raman spectra and imaging for cancerous CaCo-2 cells measured in PBS.

**Figure 8.**
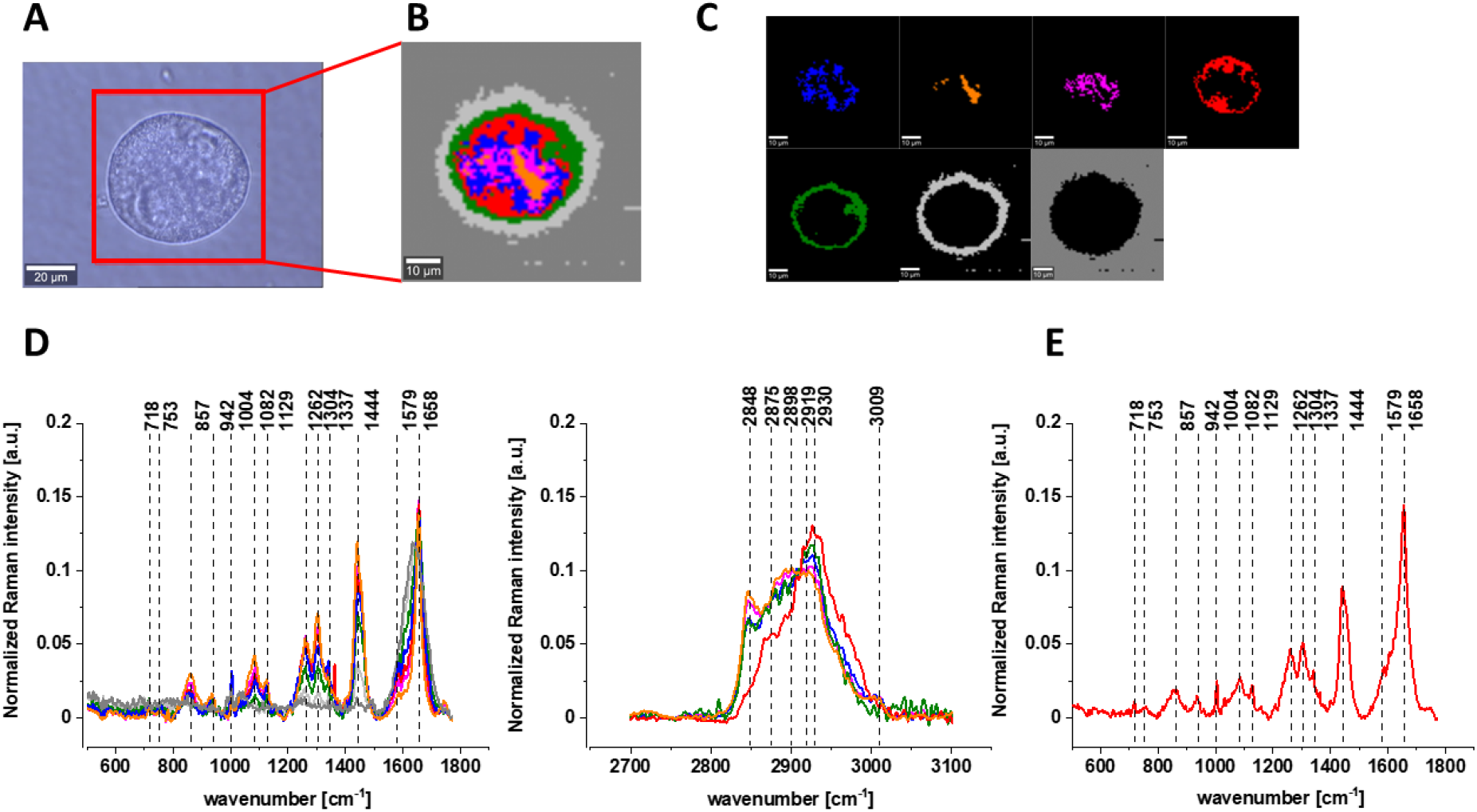
The microscopy image (A), Raman image (B) of human colon cancer single cell CaCo-2 constructed based on Cluster Analysis (CA) method, Raman images of all clusters identified by CA assigned to: lipid-rich regions (blue and orange), mitochondria (magenta), nucleus (red), cytoplasm (green), cell membrane (light grey), and cell environment (dark grey) (C), the average Raman spectra typical for CaCo-2 human cancerous colon cell for all identified clusters for low frequency and high frequency region (D), and the average Raman spectrum for human cancerous colon cell - for cell as a whole (E), all data for experiments performed without any supplementation, cells measured in PBS, colors of the spectra correspond to the colors of clusters, excitation laser line 532 nm.

Having reach this point we can start the comparative analysis of spectroscopic data.

Figures 9 shows the differential spectrum of normal and cancerous human colon cells (marked in violet color in the figure), the mean spectrum +/- SD (SD-Standard Deviation) typical for the human cancerous colon cell line (CaCo-2, marked in red), and the mean spectrum +/- SD typical for the human normal colon cell line (CCD-18 Co, marked in blue), generated using Cluster analysis (CA) for cells as a single cluster, number of cell 3.

**Figures 9.**
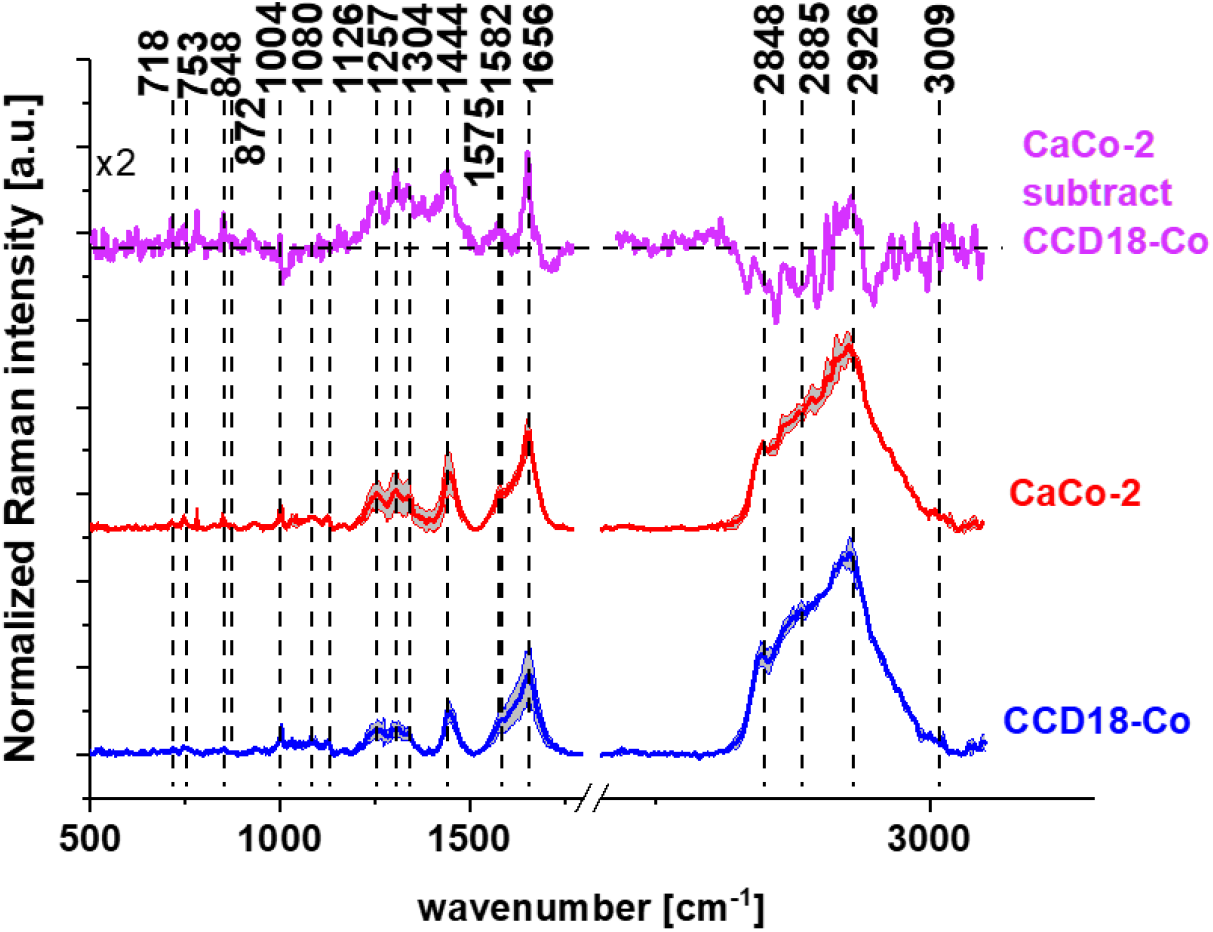
The differential spectrum of normal and cancerous human colon cells (marked in violet color), the mean spectrum +/- SD (SD-Standard Deviation) typical for the human cancerous colon cell line (CaCo-2, marked in red), and the mean spectrum +/- SD typical for the human normal colon cell line (CCD-18 Co, marked in blue), generated using Cluster analysis (CA) for cells as a single cluster, number of cell 3.

The analysis of the differential spectrum presented in Figure 9 shows that the most significant differences between normal and cancerous cells for human colon are observed for the following frequencies: 753, 872, 1082, 1575, 1004, 1585 and 2926 cm^-1^, which according to literature reports can be assigned to individual cell components such as: DNA, RNA, lipids, proteins or unsaturated fatty acids [48].

The Amide III (1230-1300 cm^-1^) and Amide I (1600-1690 cm^-1^) bands are widely used to study the secondary structure of proteins and to estimate the total amount of proteins in a test sample. The analysis of Figure 9 shows that these bands are positive on the differential spectrum, which confirms the higher amount of proteins in the biochemical composition of the Caco-2 cell line presenting cancerous cells. The bands found at 1004, 1585 and 2926 cm^-1^ are associated with proteins and also show higher intensity in cancer cells. Higher protein content in the Caco-2 cell line can be explained by the fact that cancer cells have a higher RNA/DNA content. More and more studies indicate the potential use of cell-free DNA as a biomarker of prefatory symptoms in cancer diagnosis, prognosis and monitoring. A similar relationship as for DNA and RNA can be observed for the band at 848 cm^-1^, which is attributed to mono- and disaccharides. This effect can be explained by higher concentrations of glycolysis intermediates such as acetates and lactates. All the phosphate-associated bands observed around 753, 872, 1082, 1585 cm^-1^ also show a greater proportion in the pathologically altered Caco-2 cell line. The literature has shown a higher level of phosphorylation of cancerous tissues in many organs, including the breast, brain and colon [49]. A negative correlation for the colon cancer cell line Caco-2 in Figure 9 can be observed for lipid nature high frequency bands (bands appearing in the region of 2845-2875 cm^-1^), which confirms the different lipid metabolism in normal and cancerous cells [50].

Using Raman spectroscopy, we also estimated the effect of vitamin C supplementation on the vibrational spectra of the tested cells. Figure 10 shows the Raman mean spectra determined for cells as a single cluster for CCD-18Co human normal control cells (control sample), with the addition of vitamin C (A) and the differential spectra calculated on their basis (B).

**Figure 10.**
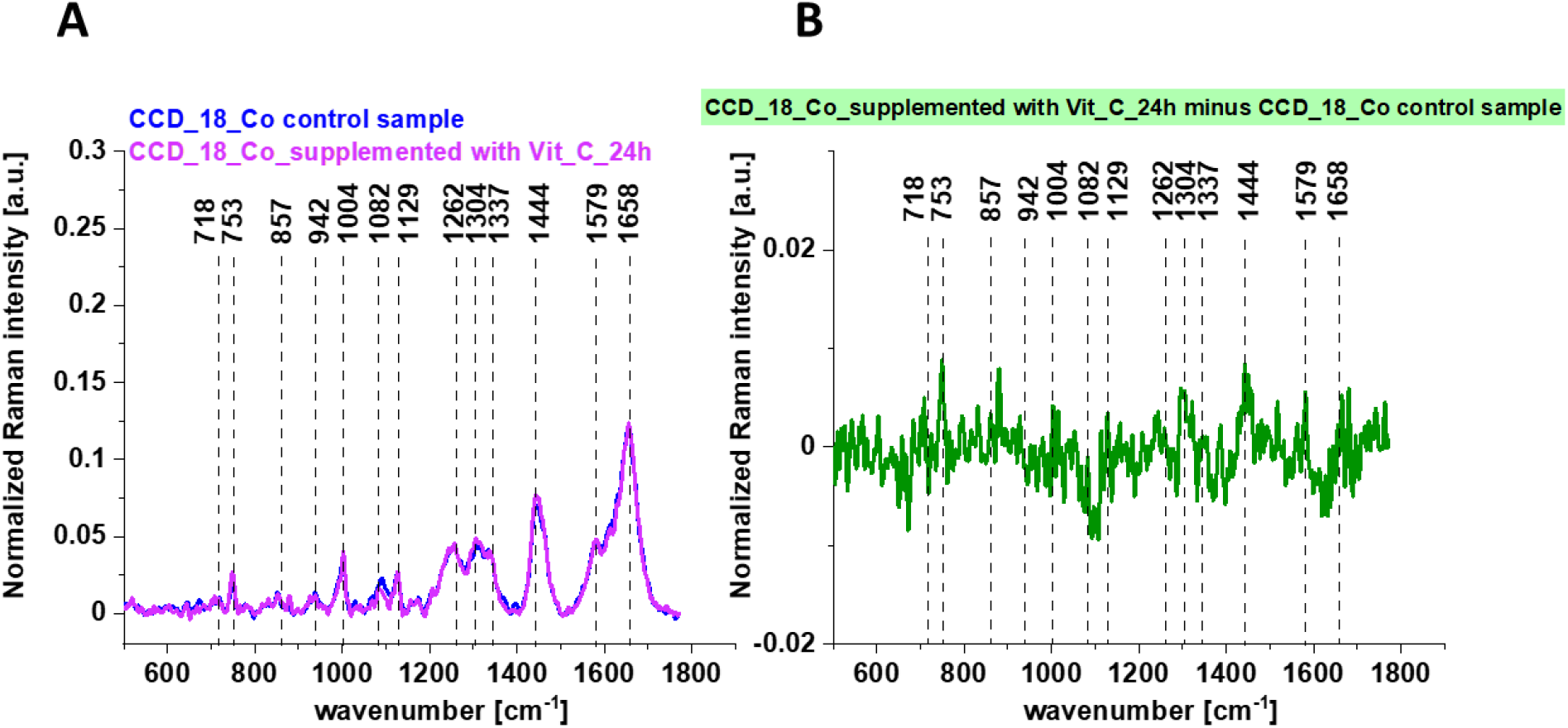
The average Raman spectra obtained for CCD-18Co human normal control cells (control sample), the average Raman spectra typical for CCD-18Co human normal control cells supplemented with vitamin C for 24h (A) and the differential Raman spectrum (B) calculated based on spectra presented on panel (A).

One can see from Figure 10 that the influence of Vitamin C on Raman spectra typical for CCD-18Co human colon normal cells is very subtle (the same result was obtained for 48h vitamin C supplementation). This observation allow to link up all differences observed for experiments performed by using tBuOOH with ROS generation and their concentration modulated by vitamin C.

Figure 11 shows the average Raman spectra of CCD-18Co human normal colon cells in oxidative stress conditions generated by using tBuOOH and for samples supplemented simultaneously by tBuOOH and vitamin C (A) and the differential spectrum calculated from them (B).

**Figure 11.**
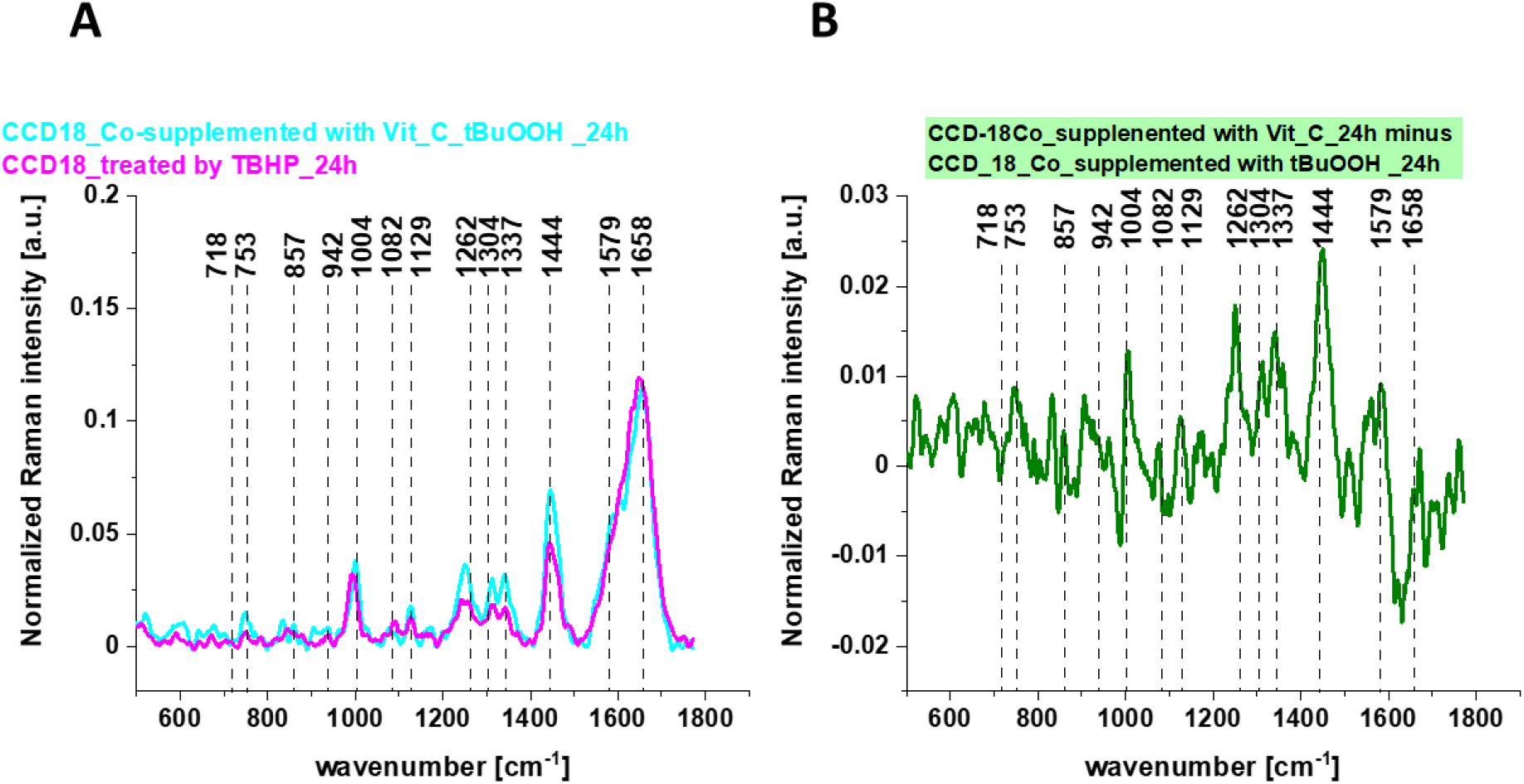
The average Raman spectra obtained for CCD-18Co human normal colon cells in oxidative stress condition, the average Raman spectra typical for CCD-18Co human normal colon cells supplemented with vitamin C and tBuOOH for 24h (A) and the differential Raman spectrum (B) calculated based on spectra presented on panel (A).

Having reached this point when the analysis based on Raman vibrational spectra for different human normal colon CCD-18Co cells groups has been performed we can compare results shown on Figures 1-8.

Considering that CCD18-Co human normal colon cells are basically composed of three types of macromolecules: proteins, nucleic acids and lipids, in order to explore characteristic changes under ROS conditions and for cells supplemented with vitamin C the qualitative and quantitative comparison between paired bands assigned to those class of compounds will be discussed according to the biological attribution of them.

Figure 12 presents an analysis based on experimental studies of the influence of oxidative stress generating factor - tBuOOH and the influence of vitamin C present in systems with generated ROS on the composition of the basic building blocks of each cell in the form of proteins, DNA and lipids. Bands 1257, 1658 cm^-1^ describe the protein content in cells, respectively: Amide I and Amide III bands, bands 750 and 1078 cm^-1^ reflect the amount of nucleic acids present in cells, bands 2854 and 3009 cm^-1^ are characteristic of lipids.

**Figure 12.**
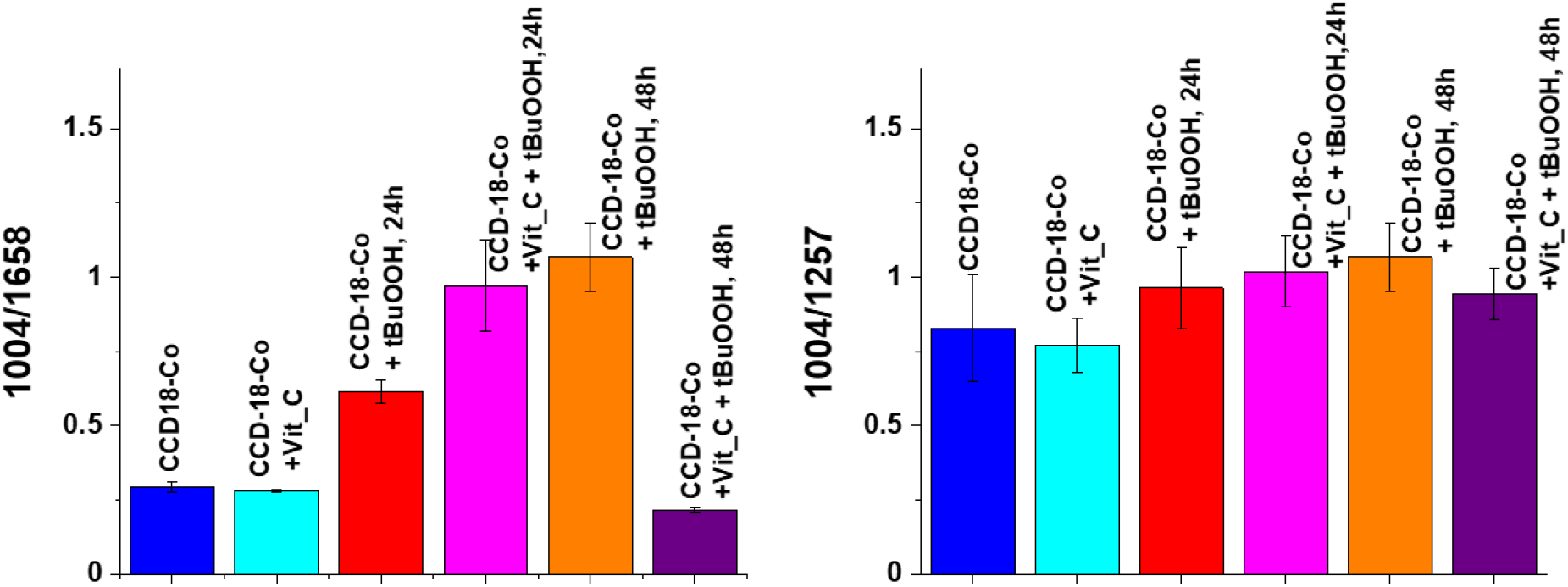
Raman band intensities rations for selected Raman bands corresponding to proteins for 6 groups of normal human colon cells CCD18-Co: control group (labeled CCD-18-Co, blue), group supplemented with vitamin C (labeled CCD-18-Co+Vit_C, turquoise), group supplemented with tBuOOH for 24h (labelled CCD-18-Co+tBuOOH, 24h, red), group. supplemented with vitamin C and tBuOOH for 24h (labelled CCD-18-Co+Vit C+tBuOOH, 24h, magenta), group supplemented with tBuOOH for 48h (labelled CCD-18-Co+tBuOOH, 48h, orange), group. supplemented with vitamin C and tBuOOH for 48h (labelled CCD-18-Co +Vit C+tBuOOH, 48h, violet).

As can be seen in Figure 12, firstly, for normal human colon CCD-18Co cells, the addition of vitamin C does not affect the protein content, while the addition of tBuOOH - ROS generating agent - modulates their amount. As the intensity of the 1004 cm^-1^ band is practically constant after the addition of vitamin C compared to non-supplemented cells, the effect of increasing the value of the ratio 1004/1658 for the CCD-18 Co + tBuOOH system results in a decrease of the intensity of the 1658 cm^-1^ band, and the modification of proteins after cell supplementation with a ROS generating compound. The results for 24 and 48 hour supplementation showed a similar tendency, although the protective effect of the antioxidant in the form of vitamin C turned out to be stronger for 48 hours.

High activity of free oxygen and nitrogen radicals in the case of proteins leads to the formation of oxidized protein products. The damage generated concerns both polypeptide chains as well as amino acid residues. The peroxidation of polypeptide chains resembles the peroxidation of lipids except for the chain nature of the observed changes. The hydroxyl radical initiates the oxidation of the protein chain by cleavage of the hydrogen atom on the α-amino acid carbon. The resulting alkyl radical reacts with oxygen to generate the product in the form of an alkylhydroperoxide. Alkylhydroperoxide can transform into an alkoxy radical, which in turn is responsible for activating the fragmentation of polypeptide chains. Breakage of the polypeptide chain can also occur as a result of free radical oxidation of aspartate, glutamate, proline. In proteins, apart from the fragmentation of the chain, as a result of ROS, oxidation of amino acid residues, including aromatic amino acid residues, may also occur. Oxidation of amino acids with a free amino, amide or hydroxyl group also leads to the formation of carbonyl derivatives present primarily in amino acid derivatives in the form of aldehydes or ketones. Modification of cysteine residues may result in breaking disulfide bonds and breaking down the tertiary structure of the protein [51].

Figure 13 presents analogous analysis performed based on Raman bands typical for nucleic acid.

**Figure 13.**
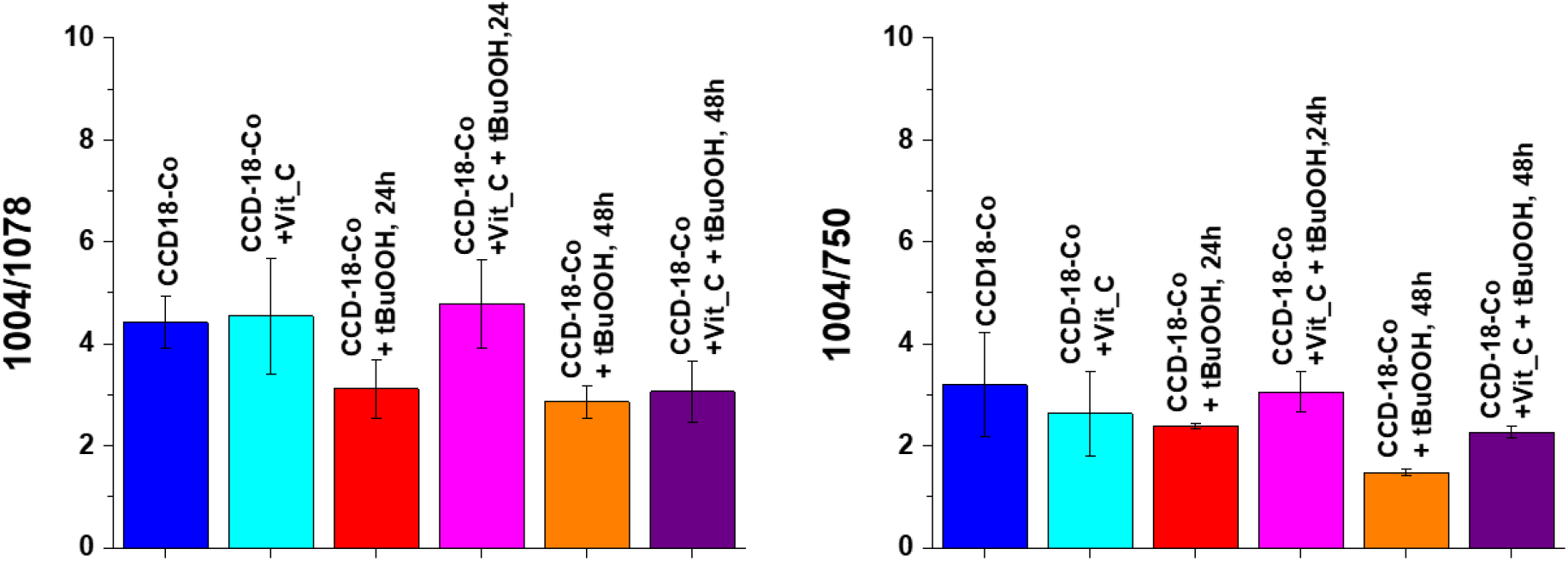
Raman band intensities rations for selected Raman bands corresponding to nucleic acids for 6 groups of normal human colon cells CCD18-Co: control group (labeled CCD-18-Co, blue), group supplemented with vitamin C (labeled CCD-18-Co+Vit_C, turquoise), group supplemented with tBuOOH for 24h (labelled CCD-18-Co+tBuOOH, 24h, red), group. supplemented with vitamin C and tBuOOH for 24h (labelled CCD-18-Co+Vit C+tBuOOH, 24h, magenta), group supplemented with tBuOOH for 48h (labelled CCD-18-Co+tBuOOH, 48h, orange), group. supplemented with vitamin C and tBuOOH for 48h (labelled CCD-18-Co +Vit C+tBuOOH, 48h, violet).

As shown in Figure 13 for normal human colon cells CCD-18Co, the addition of vitamin C does not affect the content of nucleic acids (within the standard deviation), the addition of tBuOOH - ROS generating agent, however, caused a change in the values of the analyzed Raman band intensity ratios. The addition of tBuOOH resulted in a decrease of the intensity of the bands typical of nucleic acids for both 24h and 48h of supplementation. Nucleic acids are much more stable than proteins and, when exposed to ROS, do not easily transform into a free radical state (damage is usually quickly repaired, damaged bases are “excised” and excreted from cells). Reactions of the hydroxide radical with nucleic acids damage the base sugar residues and also result in the breaking of phosphodiester bonds. Thymidine, from which thymidine peroxides are formed, is particularly susceptible to the action of radicals. The product of DNA oxygen damage is 8-hydroxy-deoxyguanosine (8-OHdG). Increased levels of 8-OHdG have been shown in the course of many diseases, including cancer, diabetes and chronic inflammation [52].

Figure 14 shows the analysis based on the bands characteristic of lipids.

**Figure 14.**
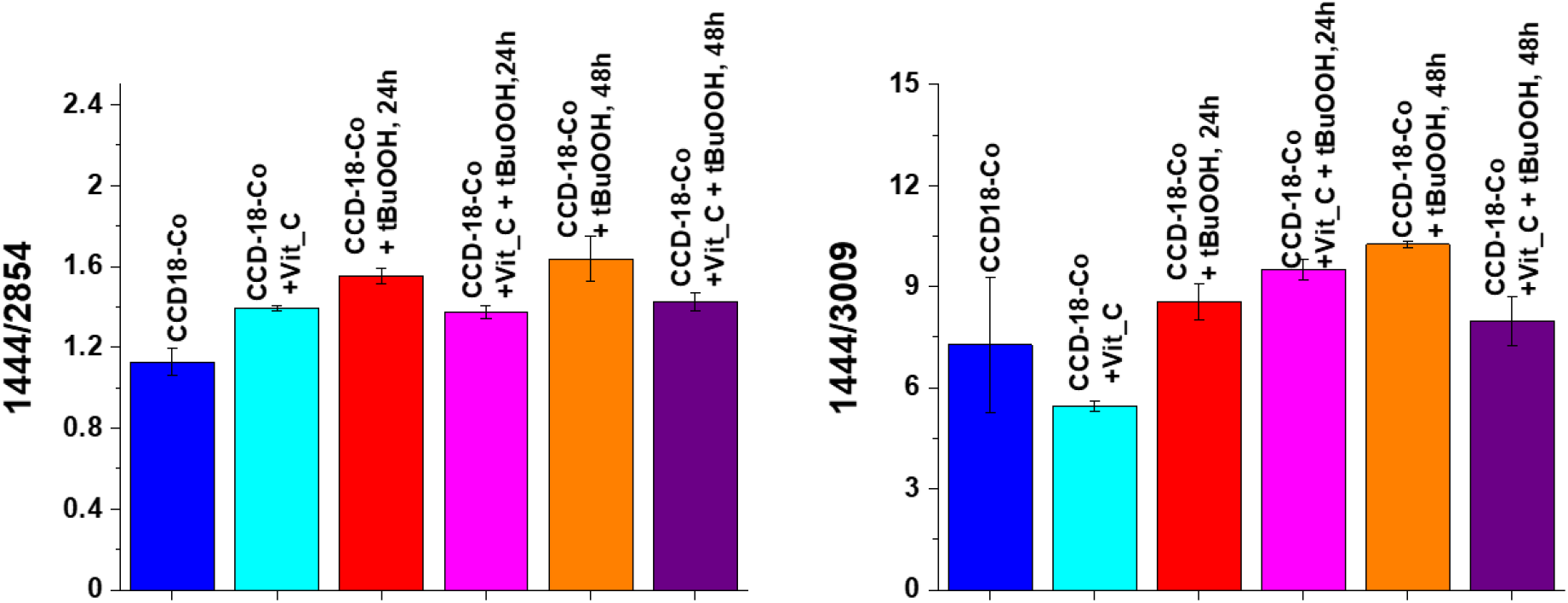
Raman band intensities rations for selected Raman bands corresponding to lipids for 6 groups of normal human colon cells CCD18-Co: control group (labeled CCD-18-Co, blue), group supplemented with vitamin C (labeled CCD-18-Co+Vit_C, turquoise), group supplemented with tBuOOH for 24h (labelled CCD-18-Co+tBuOOH, 24h, red), group. supplemented with vitamin C and tBuOOH for 24h (labelled CCD-18-Co+Vit C+tBuOOH, 24h, magenta), group supplemented with tBuOOH for 48h (labelled CCD-18-Co+tBuOOH, 48h, orange), group. supplemented with vitamin C and tBuOOH for 48h (labelled CCD-18-Co +Vit C+tBuOOH, 48h, violet).As shown in Figure 15, it also affects the lipids presence in the cells of the human colon.

Lipid peroxidation under the influence of ROS consists in the oxidation of polyunsaturated fatty acid residues, which are numerous in phospholipids, and this process takes place in a non-enzymatic way and as a result of enzymatic reactions. The product of the initiation step characteristic of lipid peroxidation are free alkyl radicals which then react with oxygen in prolongation reactions to form free peroxide radicals. These radicals located at the end of the system of double bonds react with subsequent molecules of polyunsaturated fatty acids to give peroxides (hydroperoxides) of fatty acids, which are the first products of relative stability. The termination step involves reactions directly between the radicals formed, leading to the formation of fatty acid dimers and keto or hydroxy fatty acids. The enzymatic lipid peroxidation leads to the formation of lipid hydroperoxides with the participation of enzymes from the lipoxygenases group. Enzymatic lipid peroxidation differs from non-enzymatic lipid peroxidation in that during the process with the participation of enzymes, the generated lipid peroxide radicals are converted into anions, as a result of which the substrate containing an unpaired electron is depleted, and thus the generation of hydroperoxides is inhibited. Enzymatic lipid peroxidation is also an initiating process for the formation of biologically active compounds such as prostaglandins, thromboxanes and leukotrienes. In contrast, arachidonic acid released by phospholipase A2 is the starting compound in the synthesis involving cyclooxygenases. Importantly, the products of lipid oxidation may react with proteins and DNA, contributing to their further damage [53]. The analysis of the intensity ratios of Raman bands 1444/2854 and 1444/3009 confirms the destructive effect of tBuCOOH on lipids and the “protective” nature of vitamin C both for 24 and 48h supplementation.

We have compared also the results obtained for normal human colon cells with results typical for cancer cells CaCo-2. Table 2 presents results of spectroscopic data analysis.

**Table 2.** Raman band intensities rations for selected Raman bands corresponding to proteins, nucleic acids and lipids for human normal colon cells CCD-18-Co in normal conditions, in oxidative stress conditions generated by adding tBuCOOH and for cancerous human colon cells CaCo-2.

|  | CCD-18-Co | CCD-18-Co+tBuCOOH |  | CaCo-2 |
| --- | --- | --- | --- | --- |
|  |  | 24h | 48h |  |
| **1004/1658** | 0.29±0,02 | 0,61±0,04 | 1,06±0,10 | 0.18±0.03 |
| **1004/1257** | 0.82±0,17 | 0.96±0.13 | 1.07±0.10 | 0.52±0.02 |
| **1004/1078** | 4.41±0.50 | 3.12±0.50 | 2.86±0.30 | 1.55±0.30 |
| **1004/750** | 3.19±1.00 | 2.87±0.05 | 1.47±0.07 | 1.88±0.50 |
| **1444/2854** | 1.12±0.06 | 1.55±0.03 | 1.64±0.10 | 1.49±0.04 |
| **1444/3009** | 7.27±1.99 | 8.55±0.53 | 10.25±0.08 | 8.95±1.1 |

One can see form Table 2 that intensities of Raman band ratios typical for proteins, nucleic acids and lipids depend on experimental conditions and type of human colon cells. Comparison of data obtained for normal and cancerous cells confirm that CaCo-2 cells contain more proteins and nucleic acids and less lipids (including unsaturated fraction). Comparison of data obtained for normal cells in oxidative stress conditions and cancerous type shows that the oxidation generated by adding tBuCOOH is effective but data are closer to obtained for cancerous cells CaCo-2.

## Conclusions

In the presented paper we conducted investigation on the cell culture of human colon cells using Raman spectroscopy which proves that this spectroscopic technique is a powerful tool that enables the detection of cancerous changes in biological material in an unambiguous manner. Moreover, spectroscopy and Raman imaging do not have a destructive effect on the tested biological material, so that the sample can be subjected to further analysis at a selected time.

Raman spectroscopy and imaging have been successfully applied to characterize and differentiate normal and cancerous cell line from the human colon based on the unique vibrational *‘fingerprint’* of the sample. Moreover, single cell substructures such as: the nucleus, lipid-rich regions, mitochondria, plasma membrane and cellular cytoplasm, can be accurately visualized on the basis of Raman spectra using the Cluster Analysis algorithm, which allows for grouping vibrational spectra and the creation of clusters classifying individual cell areas.

Raman studies of human colon cells show that vibrational spectroscopy can provide results that differentiate between normal and cancerous cells in the human colon based on the basic building blocks of the cells. The biochemical composition of a normal cell and a cancerous cell are different. The content of DNA, proteins and lipids, including the fraction of unsaturated fatty acids, differentiates healthy and cancerous samples. The multivariate analysis of the intensity of the lipid / protein / nucleic acid bands shows that these classes of compounds can classify the analyzed samples into normal and pathological ones.

Analysis of vibrational spectra for supplemented cells also showed that the antioxidant properties of vitamin C can be demonstrated using Raman spectroscopy and imaging based on bands characteristic of DNA, proteins and lipids.

Based on Raman band intensities attributed to proteins, nucleus acids and lipids as well as ratios 1004/1658, 1004/1257, 1004/1078, 1004/750, 1444/3009, 1444/2854 calculated based on them, we have confirmed the protective role Vitamin C for cells in oxidative stress conditions for label-free and nondestructive spectroscopic Raman method.

## Author Contributions

Conceptualization: BB-P; Funding acquisition: BB-P; Investigation: KB, BB-P; Methodology: BB-P, KB, Writing – original draft: KB, BB-P; Manuscript editing: KB, BB-P. All authors reviewed and provide feedback on the manuscripts.

## Funding

This work was supported by the National Science Centre of Poland (Narodowe Centrum Nauki) UMO-2017/25/B/ST4/01788.

## Conflicts of Interest

The authors declare no competing interests. The funders had no role in the design of the study; in the collection, analyses, or interpretation of data; in the writing of the manuscript, or in the decision to publish the results.

## Notes

### Competing Interest Statement

The authors have declared no competing interest.

